# Selenzyme: Enzyme selection tool for pathway design

**DOI:** 10.1101/188979

**Authors:** Pablo Carbonell, Jerry Wong, Neil Swainston, Eriko Takano, Nicholas J. Turner, Nigel S. Scrutton, Douglas B. Kell, Rainer Breitling, Jean-Loup Faulon

## Abstract

Synthetic biology applies the principles of engineering to biology in order to create biological functionalities not seen before in nature. One of the most exciting applications of synthetic biology is the design of new organisms with the ability to produce valuable chemicals including pharmaceuticals and biomaterials in a greener; sustainable fashion. Selecting the right enzymes to catalyze each reaction step in order to produce a desired target compound is, however, not trivial. Here, we present Selenzyme, a free online enzyme selection tool for metabolic pathway design. The user is guided through several decision steps in order to shortlist the best candidates for a given pathway step. The tool graphically presents key information about enzymes based on existing databases and tools such as: similarity of sequences and of catalyzed reactions; phylogenetic distance between source organism and intended host species; multiple alignment highlighting conserved regions, predicted catalytic site, and active regions; and relevant properties such as predicted solubility and transmembrane regions. Selenzyme provides bespoke sequence selection for automated workflows in biofoundries. The tool is integrated as part of the pathway design stage into the design-build-test-learn SYNBIOCHEM pipeline. The Selenzyme web server is available at http://selenzyme.synbiochem.co.uk.

## Introduction

In recent years, powerful bioinformatics tools are being increasingly integrated into systems and synthetic biology pipelines (Kell, 2006; Carbonell *et al.*, 2016). Synthetic biology employs the engineering principle of an iterative Design-Build-Test-Learn cycle. In the case of developing engineered organisms for the production of high-value compounds, the Design stage involves the identification of the most suitable combinations of starting substrates, enzymes, regulatory components and chassis organism for the desired biosynthetic pathway. Hence, bioinformatics tools used at this stage usually carry out database mining in order to search for the best candidate parts and devices. Tools such as BNICE (Hadadi *et al.*, 2016) or RetroPath (Carbonell *et al.*, 2014) are capable of establishing possible pathways leading to a target compound by using generalized reaction rules. RetroPath also provides an initial assessment of pathway feasibility by ranking candidate enzyme sequences in an automated way using machine learning (Carbonell *et al.*, 2011). In order to improve the ability to select candidate sequences beyond purely automated selection, some more specialized tools are available, including antiSMASH for biosynthetic gene clusters (Weber *et al.*, 2015), CanOE for orphan reactions (Smith *et al.*, 2012), as well as tools based on reaction homologies like MRE (Kuwahara *et al.*, 2016) and EC-Blast (Rahman *et al.*, 2014) or machine learning (Mellor *et al.*, 2016). Here, we extend such capabilities through Selenzyme, a sequence selection with the ability to mine generalized reaction rules that is integrated into a larger computational design pipeline for metabolic engineering within the SYNBIOCHEM Centre that includes the RetroPath 2.0 workflow (Delepine *et al.*, 2017) and will be integrated with the downstream tool PartsGenie (Swainston, Dunstan, *et al.*, 2017). In this way, this new tool can assist in the selection of heterologous candidate enzymes to express for both natural and non-natural targets.

## Design and implementation

Our goal is to mine for candidate enzyme sequences for any desired target reaction or set of reactions in a pathway. A unique feature of Selenzyme is that target reactions do not necessarily need to exist in databases, enabling the tool to search for alternative routes for biosynthesis, degradation and transport either for natural or non-natural products.

## Data sources

Selenzyme uses SYNBIOCHEM’s biochem4j graph database (Swainston, Batista-Navarro, *et al.*, 2017) (June 2017) as its main data source. The database contains relevant information on known relationships between reactions (36765), chemicals (19735), enzymes (245704) and organisms (8431).

## Reaction screening

A Selenzyme query consists of a target reaction, represented in.rxn format (MDL) or a character string using either the SMARTS format or the derivative SMIRKS representation for generalized reactions (Daylight Theory Manual, Version 4.9. http://www.daylight.com/ (accessed 1 September 2017)). The query is then screened against the reaction database in order to look for similar chemical transformations. Reaction similarity is computed by first determining the most similar reactant on each side of the reaction for each substrate and each product. Arithmetic mean similarities for subsets (substrate, left reactant), (product, right reactant) and (substrate, right reactant), (product, left reactant) are computed, and the resulting overall similarity for each pair is obtained through the application of the quadratic mean (RMS) to give higher weight to larger similarities. Individual Tanimoto similarities between chemicals are obtained using preselected fingerprints, i.e., circular (Morgan), based on molecular subgraphs (RDKit) or optimized for substructure screening (patterned) (Landrum, 2017). This algorithm provides a fast way for calculating reaction similarities that, in our experience, achieves a good trade-off between reaction specificity and calculation speed. Similarity between the query reaction and the database is calculated both in forward and reverse directions as often reaction directionality in the database is unknown or has not been appropriately annotated. The user can choose to rank similarities in both directions or to use only the direction of the reaction based on a consensus list that has been generated according to reaction information based on curated information from MetaCyc (see supplementary material for a description of the procedure).

## Output format

The algorithm proceeds through the list of reactions ranked by decreasing similarity and collects annotated sequences associated with the reactions down to the desired number of targets, which is a user-configurable parameter. For each sequence, several useful properties such as source organism, phylogenetic distance to the host chassis, predicted secondary structure and physicochemical properties computed via the EMBOSS suite are collected. Optionally, a multiple sequence alignment (MSA) can be generated using T-Coffee (Taly *et al.*, 2011). The output generates a.csv file containing the list of top sequence candidates initially ranked based on reaction similarity followed by conservation score (if the MSA is available) and phylogenetic distance, along with associated properties and references, as well as a fasta file with the Uniprot sequences.

## Web server and RESTful service

On top of this core tool, there is a web server built using Flask in Python that provides a web interface where the user can input the reaction query as an.rxn file or a SMARTS string. Once the reaction format has been verified, the query is submitted and the ranked list of sequence candidates is presented as an interactive table, which can be sorted on user-defined summary scores based on a weighted average of selected columns or properties. Additional sequences can be added or removed from the table. The MSA can be visualized through MSAviewer (Yachdav *et al.*, 2016). The SMARTS query can be submitted through a GET request as well, allowing in this way to easily generate query links pointing to the Selenzyme web service.

Moreover, a RESTful service has been implemented, so that Selenzyme can accept multiple queries from any other web-based application. As an example of the application of the REST service, a KNIME node (O’Hagan and Kell, 2015; Fillbrunn *et al.*, 2017) is available so that the reaction query can be generated from chemoinformatics operations within a workflow (examples available at http://www.myexperiment.org/packs/734, see supplementary information). Similarly, the resulting output tables containing the sequences can easily be processed downstream in the workflow or sent to other sequence analysis services such as Galaxy.

## Guided example

Selenzyme is a flexible tool that can provide enzyme selection solutions in different scenarios. One typical application case is in the selection of enzyme sequences to engineer a biosynthetic pathway. We consider for instance the biosynthetic pathway of the flavonoid pinocembrin engineered in the chassis organism *Escherichia coli*. The pathway consists of 4 reaction steps (PAL, 4CL, CHS, and CHI). The reactions were defined using a molecule editor and exported as SMILES strings. Each string was manually submitted to Selenzyme in SMILES format. Table 1 shows the 10 top selected sequences for PAL based on the predefined combined score of reaction similarity, phylogenetic distance and sequence conservation.

**Figure 1.**
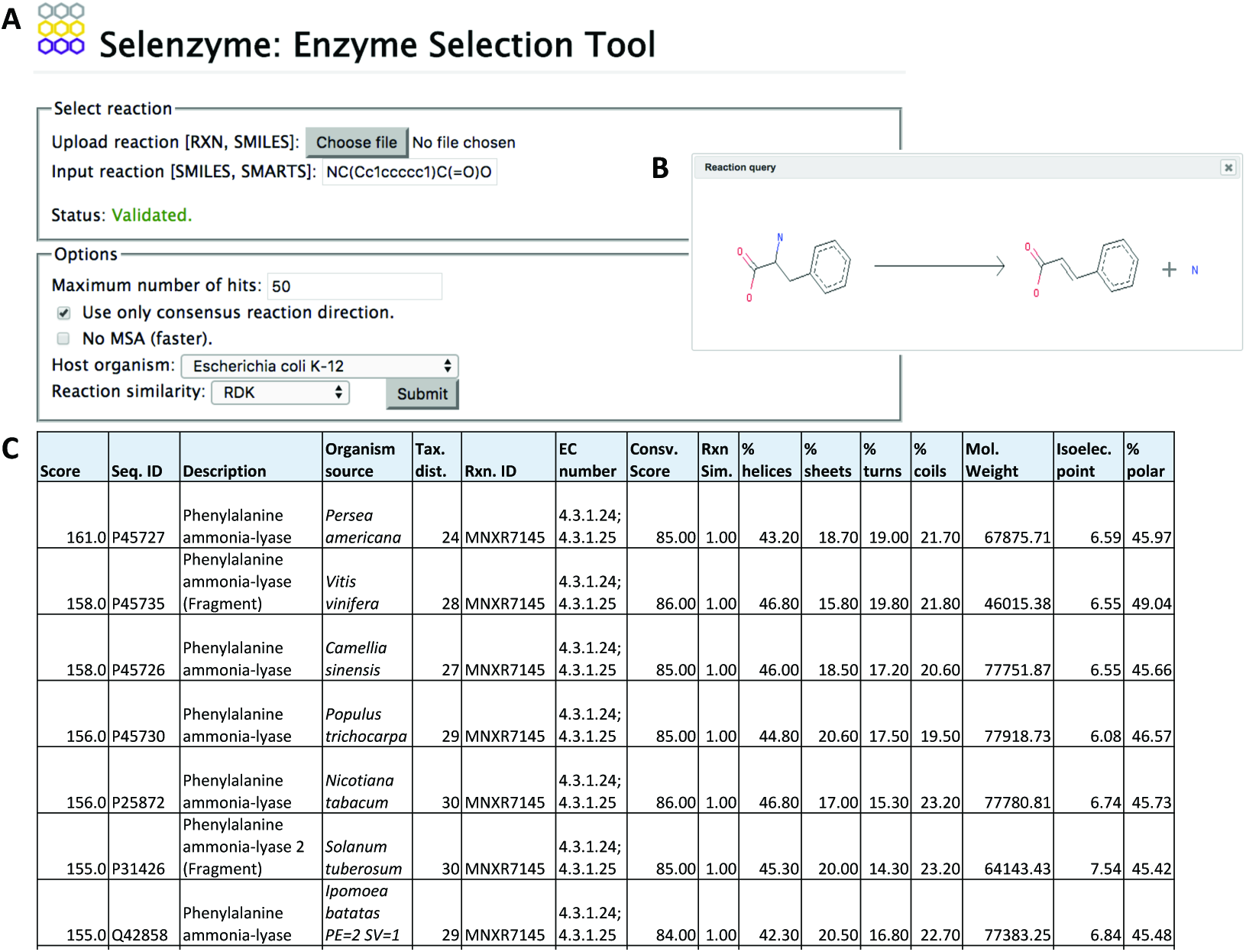
Query submission to Selenzyme through the web interface and resulting table of top ranked sequences on the default score with associated properties and cross-references to databases. (**A**) Only reactions in the preferred direction in the database were considered for the target reaction (**B**). (**C**) The results page provides links for downloading the table, or the multiple sequence alignment, as well as links to external databases.

